# The Autoimmune Susceptibility Gene, *PTPN2*, Restricts Expansion of a Novel Mouse Adherent-Invasive *E. coli*

**DOI:** 10.1101/709634

**Authors:** Ali Shawki, Rocio Alvarez, Marianne R. Spalinger, Paul M. Ruegger, Anica Sayoc, Alina N. Santos, Pritha Chatterjee, Jonathan Mitchell, John Macbeth, Michel L. Tremblay, Ansel Hsiao, James Borneman, Declan F. McCole

## Abstract

Inflammatory bowel diseases (IBD) involve genetic and environmental factors that play major roles in disease pathogenesis. Loss-of-function single-nucleotide polymorphisms (SNPs) in the protein tyrosine phosphatase non-receptor type 2 (*PTPN2*) gene increase the risk of IBD and are associated with altered microbiome population dynamics in IBD. Moreover, expansion of intestinal pathobionts, such as adherent-invasive *E. coli* (AIEC), is strongly implicated in the pathogenesis of IBD as AIEC increases pro-inflammatory cytokine production and alters tight junction protein regulation suggesting a potential mechanism of pathogen-induced barrier dysfunction and inflammation. The aim of this study was to identify if PTPN2 deficiency disturbs the composition of the intestinal microbiome to promote expansion of specific bacteria with pathogenic properties. In mice constitutively lacking *Ptpn2* we identified increased abundance of a novel adherent-invasive *E. coli* (AIEC) that showed similar adherence and invasion of intestinal epithelial cells, but greater survival in macrophages to the IBD associated AIEC, LF82. Furthermore, we confirmed this novel mouse AIEC (*m*AIEC) caused disease when administered to germ-free and mice lacking segmented-filamentous bacteria (SFB). Moreover, *m*AIEC infection increased severity of and prevented recovery from dextran-sodium sulfate (DSS)-induced colitis. *m*AIEC genome sequence analysis showed >90% similarity to LF82. Interestingly, *m*AIEC contained distinct attachment genes not found in LF82 thereby also demonstrating the novelty of this AIEC. We show here for the first time that an IBD susceptibility gene, *PTPN2*, plays a key role in modulating the gut microbiome to protect against a novel pathobiont. This study generates new insights into gene-environment-microbiome interactions in IBD.

## Introduction

Genome-Wide Association Studies (GWAS) have identified an association between loss-of-function single-nucleotide polymorphisms (SNPs) in the protein tyrosine phosphatase non-receptor type 2 (*PTPN2*) gene, that encodes the T-cell protein tyrosine phosphatase (TCPTP), and several autoimmune diseases including Crohn’s disease, ulcerative colitis, celiac disease, type 1 diabetes and rheumatoid arthritis (Jostins et al., 2012; Franke et al., 2008; Wellcome Trust Case Control Study, 2007; Spalinger et al., 2016b; Smyth et al., 2008). Crohn’s disease (CD) and ulcerative colitis (UC), collectively known as inflammatory bowel disease (IBD), are chronic intestinal inflammatory conditions whose etiology is unclear. Several factors such as genetics and alterations in the microbiome are critical determinants of IBD pathogenesis (Khor et al., 2011). PTPN2 (TCPTP) has an essential role in restricting inflammation as homozygous *Ptpn2* knockout mice exhibit substantially increased expression of pro-inflammatory cytokines and uncontrolled systemic inflammation (You-Ten et al., 1997; Heinonen et al., 2004). TCPTP regulation of inflammation is due, at least in part, to restriction of pro-inflammatory signaling pathways mediated by members of the Janus kinases (JAK) and signal transducer and activator of transcription (STAT) families of signaling kinases (JAK-STATs), which are activated by inflammatory cytokines such as IFN-γ (Yamamoto et al., 2002; Heinonen et al., 2004).

With respect to intestinal inflammation, TCPTP restricts intestinal epithelial cell (IEC) barrier defects and tight junction remodeling induced by IFN-γ, an inflammatory cytokine involved in several autoimmune diseases including IBD and celiac disease (Yamamoto et al., 2002; Heinonen et al., 2004; Krishnan and McCole, 2017; Scharl et al., 2009). The intestinal epithelium forms a selectively permeable barrier between the lumen and the sub-mucosa through the formation of multiprotein complexes of desmosomes, adherens junctions, and tight junctions (TJ) that regulate paracellular permeability (Matter and Balda, 2003; Shen et al., 2011). The intestinal epithelial barrier is essential to maintain appropriate compartmentalization of tissue vs. lumenal factors in the gut. Specifically, the epithelium ensures that lumenal microbes and their products are restricted from accessing lamina propria immune cells or gaining access to the underlying vasculature. Indeed, when intestinal epithelial integrity is compromised translocation of bacteria, and bacterial products such as lipopolysaccharide, can occur and trigger inflammatory responses that in severe cases lead to sepsis (Assimakopoulos et al., 2018; Guerville and Boudry, 2016; Bein et al., 2017).

Approximately 4 × 10^13^ bacteria exist in the gastrointestinal tract with more than 35,000 bacterial species that play a major role in maintaining intestinal homeostasis (Chowdhury et al., 2007; Ghoshal et al., 2012; Sender et al., 2016). Alterations in the intestinal microbiome are a major environmental factor in the pathogenesis of IBD (Stecher, 2015; Rogler et al., 2018; Craven et al., 2012). Expansion of pathobionts, such as adherent-invasive *Escherichia coli* (AIEC), is associated with IBD pathogenesis. This is likely due to a combination of pathological activities of AIEC which include induction of pro-inflammatory cytokine (IFN-γ, TNF-α, IL-13) production; increasing susceptibility to induced colitis in a genetically-susceptible host; and disrupting expression and distribution of epithelial TJ proteins leading to increased intestinal permeability (Martinez-Medina and Garcia-Gil, 2014; Chassaing et al., 2014; Carvalho et al., 2012; Darfeuille-Michaud et al., 2004; Eaves-Pyles et al., 2008; Berkes et al., 2003; Shawki and McCole, 2017; Liao et al., 2008; Ulluwishewa et al., 2011).

A major gap in our understanding of how complex inflammatory diseases such as IBD arise, relates to how genetic susceptibility alters the intestinal environmental to favor the expansion of commensal microbes with pathogenic potential prior to manifestation of clinical disease (Molodecky and Kaplan, 2010). Clinical genetic studies have provided some clues in this regard. A role for *PTPN2* as a clinically relevant modulator of the gut microbiome was first identified in IBD patients where *PTPN2* SNPs influenced the population dynamics of intestinal microbes, while a separate study also identified dysbiosis and increased disease severity in patients harboring *PTPN2* SNPs (Knights et al., 2014; Yilmaz et al., 2018). 16S analysis showed that *Ptpn2* deficiency in mouse CD4 T-lymphocytes aggravated adoptive T-cell transfer colitis and promoted a broad dysbiosis featuring typical colitis-associated increases in *Bacteroidetes* and *Proteobacteria*, and decreased *Firmicutes*, in stool (Spalinger et al., 2015). In this study we aimed to use a more rigorous approach of sequencing the internal transcribed spacer (ITS) variable region of bacteria to identify if constitutive TCPTP loss alters intestinal microbiome composition and identify specific bacterial species modulated by *PTPN2*.

Here, we report that *PTPN2*-deficient mice exhibit a highly significant increase in abundance of a novel mouse adherent-invasive *E. coli* that has significant, but also distinct, genetic overlap with the human LF82 isolated from Crohn’s disease patients. This mouse AIEC (*m*AIEC) caused weight loss in germ-free mice and both exacerbated and delayed recovery from inflammatory colitis thus confirming its pathogenic properties. Our data have identified a novel mouse AIEC that may prove a valuable tool for the study of pathobiont-induced disease and for the interactions of genetic and environmental factors contributing to autoimmune diseases such as IBD.

## Methods

### Animal Procedures

#### Ethical Statement on Mouse Studies

All animal care and procedures were performed in accordance with institutional guidelines and approved by the University of California Riverside Institutional Animal Care and Use Committee under Protocol# A20190001B.

#### Housing and Husbandry of Experimental Animals

Constitutive *Ptpn2* knockout (KO) male and female mice were generated by breeding of heterozygous (Het) mice on a BALB/c background and genotyped as previously described (You-Ten et al., 1997). Wild-Type (WT) and *Ptpn2*-Het littermate male and female mice were used as controls. All mice used for microbiome analysis were approximately 3 weeks old (19-23 days of age) at time of sacrifice and were housed in a specific pathogen-free (SPF) vivarium at UC Riverside.

Wild-type 10-week old confirmed SFB-free C57Bl/6 female mice were purchased from JAX labs (Stock# 000664) and housed in a SPF vivarium.

Germ-free ∼12-week old C57Bl/6 male and female mice generated from within the germ-free facility at UC Riverside were kindly provided by Dr. Ansel Hsiao.

#### Microbiome studies

Lumenal contents (distal ileum, cecum, proximal and distal colon) and mucosal-associated microbes (small and large intestine) were isolated as previously published (Presley et al., 2010). Briefly, DNA was extracted using the PowerSoil DNA Isolation Kit (MO BIO Laboratories, Carlsbad, CA, USA), and a 30-second beat-beating step using a Mini-Beadbeater-16 (BioSpec Products, Bartlesville, OK, USA). Bacterial ribosomal RNA (rRNA) internal transcribed spacer (ITS) region sequencing (250 base reads) using the Illumina MiSeq platform was performed as previously described (Ruegger et al., 2014).

#### Bacterial isolation and culture

Mouse AIEC (*m*AIEC) was isolated from fecal content from a *Ptpn2*-KO mouse, plated onto *E. coli* media (ChromoSelect Agar B; Sigma Aldrich) per manufacturer protocol. Bacterial colonies with 100% sequence match to a 250bp ITS segment to the human AIEC LF82 were subcultured to obtain pure *m*AIEC cultures. Bacteria from stocks frozen at –80°C in 1:1 glycerol:LB were cultured overnight in Luria–Bertani (LB) broth at 37°C, 225rpm, and regrown the next day in fresh LB to exponential phase growth. Culture was pelleted, washed with phosphate buffered saline (PBS), and resuspended in PBS.

#### Bacterial infection studies

The bacteria used were the LF82 human AIEC kindly provided by Dr. Arlette Darfeuille-Michaud, *m*AIEC, and K12 (a non-invasive *E. coli*, ATCC 25404). All cells were seeded at 2 × 10^5^ cells per well in 24-well plates, cultured until confluent, and infected at a multiplicity of infection (MOI) of 10 bacteria/cell as previously described (Martinez-Medina et al., 2009; Baumgart et al., 2007). Media was changed to antibiotic-free media 24 hrs prior to infection. For adherence and invasion studies, Caco-2 brush border (Caco-2bbe) intestinal epithelial cells (IECs) were infected with bacteria for 3 hrs. Bacterial survival was performed using murine J774 macrophages (Mϕ) infected with bacteria for 2 hrs followed by a PBS wash, incubation with fresh media containing 100µg/mL Gentamicin for 1 hr followed by incubation with fresh media containing 20µg/mL Gentamicin for 24 hrs. After bacterial infection, cells were washed with PBS and lysed with 1% Triton-X for 5 min. Lysates were plated onto Luria Bertani Agar plates and cultured overnight at 37°C.

Mice were infected by oral gavage with 10^9^ colony forming units (CFU)/mL of *m*AIEC or K12 in 100µL PBS/mouse. Body weight was monitored daily. Colonization and translocation was measured by overnight culture of homogenized tissue suspended in 500µL of PBS.

#### Induction and assessment of colitis severity and histological score

Colitis was induced by supplementation of drinking water with dextran-sodium sulfate (DSS) at 2.5% for 7 days as described (Kasper et al., 2016). Mice were randomized to three groups: 1) <u>H_2_O group</u> did not receive DSS but received daily single gavages of PBS (H_2_O-PBS) or bacteria (H_2_O-K12, H_2_O-*m*AIEC) for 4 consecutive days; 2) <u>acute DSS group</u> received DSS for 7 days with daily single gavages of PBS (DSS-PBS) or bacteria (DSS-K12, DSS-*m*AIEC) for the first 4 consecutive days of DSS treatment followed by 4 days of recovery post DSS treatment; and 3) <u>recovery from colitis group</u> received DSS for 7 days with daily single gavages of PBS (rec-PBS) or bacteria (rec-K12, rec-*m*AIEC) for 4 consecutive days post DSS treatment.

To assess disease severity of colitis in animals, disease activity index (DAI) was monitored daily (Becker et al., 2006; Friedman et al., 2009). Myeloperoxidase, an indicator of infiltration/activation of neutrophilic granulocytes was measured as previously described (Spalinger et al., 2016a; Friedman et al., 2009). Histological scoring for inflammatory infiltration and epithelial cell damage was performed on H&E-stained sections of the most distal 1 cm of the mouse colon (Becker et al., 2006; Spalinger et al., 2015; Friedman et al., 2009). Briefly, sections from the most distal 1 cm of the mouse colon were deparaffinized in Citrisolv and dehydrated using a series of alcohol washes with decreasing concentration, before staining with Hematoxylin for 10 min. The sections were then rinsed with tap water for 10 mins and stained in Eosin Y solution for 15 sec, briefly rinsed in Milli-Q H_2_O and dehydrated in ethanol with ascending concentrations followed by Citrisolv, and finally mounted with Permount (ThermoFisher Scientific, Waltham, MA). Microscopic assessment was performed using a Leica DM5500 microscope attached to a DFC365 FX camera using a 63× oil immersion objective with an additional 2× digital zoom. The individual images were converted to tiff files with the LAS-V4.12 lite software, and Photoshop (Adobe) was used to create the final figures.

#### In-vivo barrier permeability

Mice were gavaged with fluorescein isothiocyanate (FITC)-dextran (4 kDa, 80 mg/mL). After 5 hrs, blood was collected by retro-orbital bleed into serum collection tubes. The blood was centrifuged at 4°C, 8’000g, for 3 min, and serum analyzed for FITC-dextran concentration with the Veritas Microplate luminometer (Turner Biosystems, Sunnyvale, CA), GloMax software (Promega, Madison, WI), using an excitation wavelength of 485 nm and an emission wavelength of 535 nm. Standard curves for calculating FITC-dextran concentration in the samples were obtained by diluting FITC-dextran in water.

#### Genome sequence analysis

Bacteria were cultured and pelleted as described above. Samples were sent to Novogene (Sacramento, CA) for DNA isolation, library construction, and sequencing of bacterial genomes. The genome of *m*AIEC was compared with the genome sequence of the human LF82 isolate (Darfeuille-Michaud et al., 1998). Putative attachment genes found in *E. coli’s* and other AIECs including LF82 were identified by comparing the known sequence of each gene from K12. Attachment genes common between *m*AIEC, another non-pathogenic mouse AIEC NC101, and the human AIEC LF82 or unique to each isolate were compared and percent similarity of each gene between each isolate was determined (Kim et al., 2005).

#### Statistical analyses

We set critical significance level α = 0.05 and analyzed our data using parametric statistics. Data are expressed as mean, SD for *n* independent observations per group unless stated otherwise. Between-group inferences were made by using one-way or 2-way analysis of variance (ANOVA) and controlled by the false discovery rate (FDR) procedure where applicable (Curran-Everett, 2000). Microbiome analyses were performed as previously described (Ginnan et al., 2018).

## Results

### Constitutive Ptpn2-deficient Mice Have an Altered Intestinal Microbiome with Expansion of Proteobacteria

Since *PTPN2* genotyped human IBD patients show altered intestinal microbiomes, we analyzed the microbiomes of *Ptpn2*-deficient mice (Yilmaz et al., 2018). Constitutive KO mice show distinct lumenal bacterial communities compared with WT and Het mice (Figure 1A), and this was not dependent on intestinal region since there was no difference between lumenal bacterial communities in cecum, proximal colon, and distal colon (Figure 1B). When phyla levels were examined in littermates from the same cages, the major bacterial phylotypes were *Bacteroidetes, Firmicutes*, and *Proteobacteria* in WT mice (Figure 1C). However, KO mice, and to a lesser extent, Het mice, show reduced abundance of *Bacteroidetes* compared with WT mice. *Proteobacteria* was the phylum with the greatest increase in KO compared with WT and Het mice (Figure 1C). These data showed that loss of *Ptpn2* induced a shift in the microbiome that was not dependent on large intestinal region.

**Figure 1.**
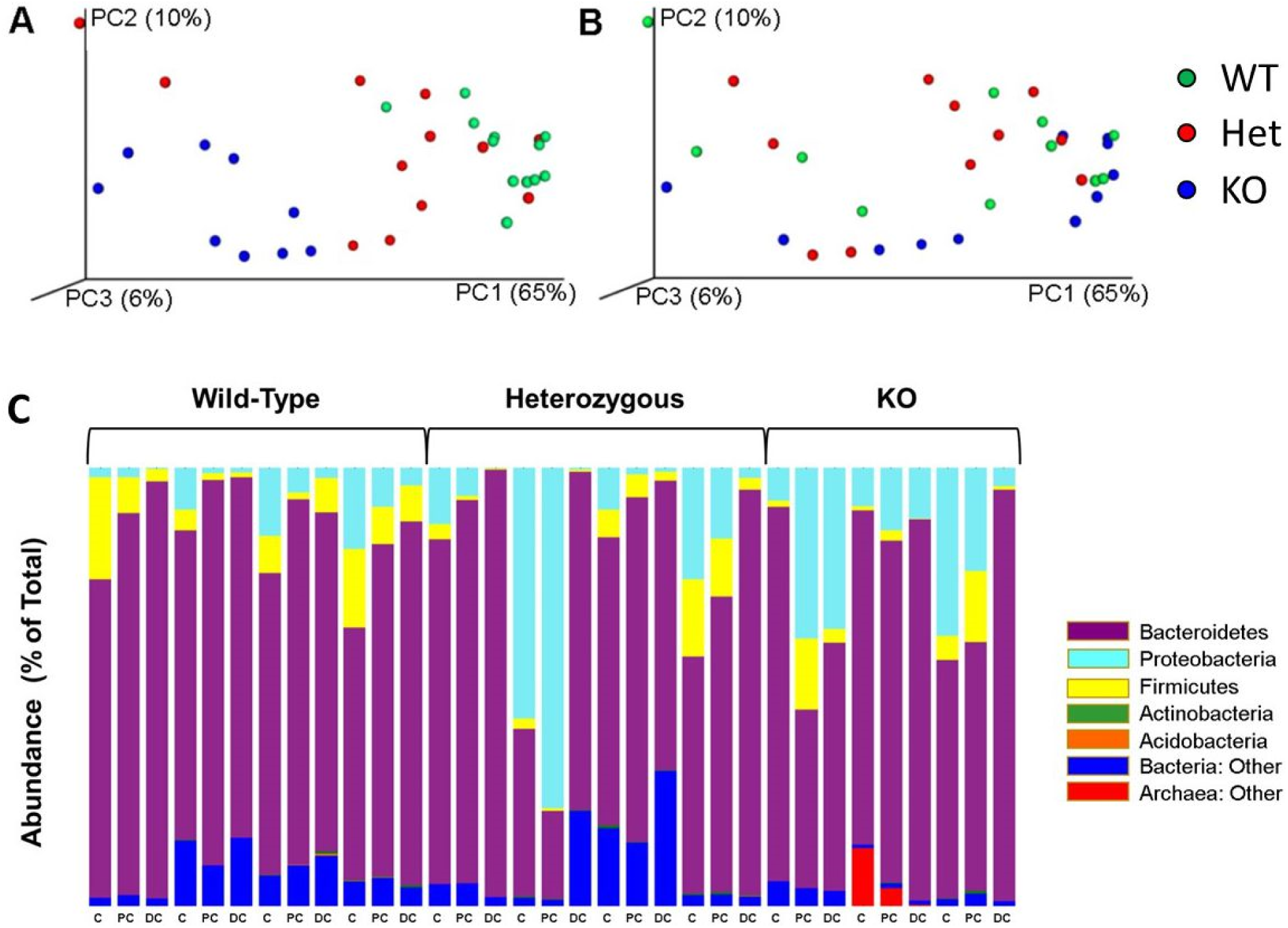
Lumenal bacterial communities display greater variance by *Ptpn2* mouse genotype than intestinal region. Lumenal bacterial communities represented as single points in a three-dimension plot with the percent values of each axis indicating the amount of variance in the data that is explained by that axis. ***A***, Depiction of the three different *Ptpn2* genotypes: WT (green), Het (red) and KO (blue). ***B***, Depiction of the three different intestinal regions: cecum (red), proximal (green) and distal (blue) colon. These comparisons used principal coordinates analysis of weighted UniFrac values (Lozupone and Knight, 2005). For each genotype, samples from the three intestinal regions were pooled. For each intestinal region, samples from the three genotypes were pooled. Percent values of each axis indicates the variance (*P*=0.001 for all 3 pairs; Adonis test). ***C***, Differentially abundant bacterial phylotypes in three different intestinal regions (C = cecum; PC = proximal colon; DC = distal colon) of mice with three different *Ptpn2* genotypes (Wild-Type, Heterozygous, Knockout (KO). Values are % of rRNA ITS sequencing reads.

### Ptpn2-deficient Mice Display Significant Expansion of an IBD-Relevant Adherent-Invasive E. coli

To further characterize the phylum *Proteobacteria*, which included *Enterobacteria* and *Escherichia coli* (*E. coli*), we probed for strains highly abundant within this phylum in lumenal and mucosal associated content from small (ileum) and large intestine. Consistent with the overall increase in abundance of *Proteobacteria* (Figure 1C), the *E. coli* phylotype had greater than 2% of the total sequencing reads and its abundance was increased to 7.3% in KO mice compared with WT (5.4 × 10^−5^%) and Het (1.1 × 10^−4^%) mice (Figure 2A). Interestingly, there was 100% sequence identity and 100% sequence coverage of a 250 base pair internal transcribed spacer (ITS) region with 6 sequences in the nt database (GenBank), and 2 of the 6 were AIECs (LF82, accession CU651637; O83:H1 strain NRG 857C, accession CP001855). We tentatively assigned this novel *E. coli* as mouse AIEC (*m*AIEC).

**Figure 2.**
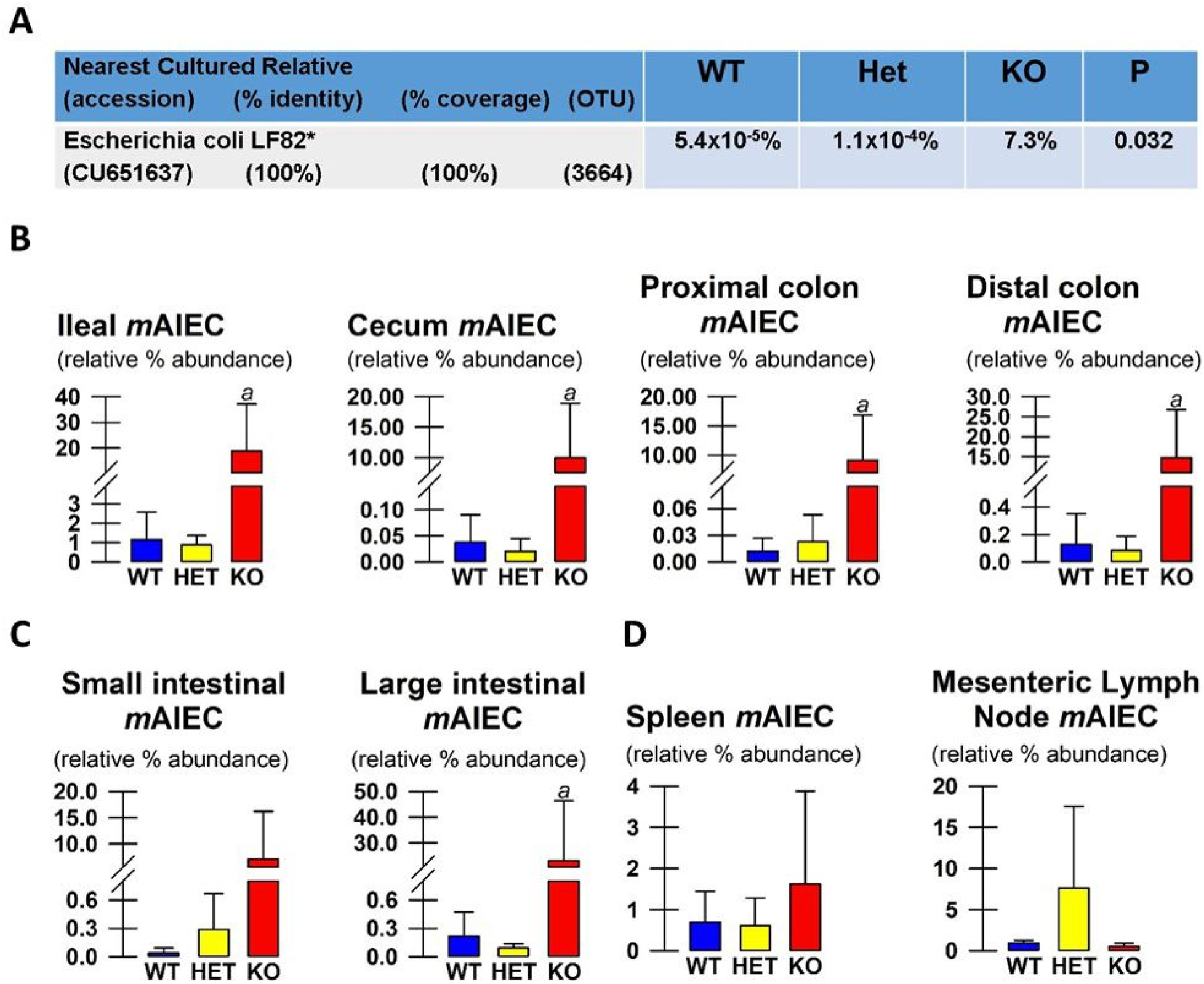
Expansion of an IBD-relevant *E. coli*. To further characterize the phylum *Proteobacteria*, which includes *Enterobacteria* and *E. coli*, we probed for strains highly abundant within this phylum in lumenal and mucosal-associated bacteria. ***A***, Consistent with the overall increase in abundance of *Proteobacteria* in Fig 1C, the *E. coli* phylotype had greater than 2% of the total sequencing reads and 100% identity and 100% coverage with 6 sequences in the nt database (GenBank), and 2 of the 6 were AIECs (LF82, accession CU651637 ; O83:H1 strain NRG 857C, accession CP001855). Values are % of rRNA ITS sequencing reads. % identity and % coverage were determined by analyses using Blast (NCBI) (Altschul et al., 1990). Abundance values were compared using ANOVA (*n* = 5-8 mice for each of the three genotypes). ***B***, Lumenal (*n* = 5-8) and ***C***, mucosal-associated *m*AIEC (*n* = 5-8) and translocation of *m*AIEC into the ***D***, spleen and mesenteric lymph node (*n* = 3-4) from Wild-Type (WT) and *Ptpn2* Heterozygous (Het) and Knockout (KO) mice. Values are % of rRNA ITS sequencing reads within the *Proteobacteria* phylum. One-way ANOVA revealed an interaction in lumenal (*P* ≤ 0.02) and mucosal regions tested (*P* ≤ 0.03) except in small intestine (*P* = 0.06) and no interaction in spleen or MLN *m*AIEC (*P* ≥ 0.19). Post-hoc analysis of significant interactions revealed higher *m*AIEC abundance in KO mice compared with WT and Het mice (^*a*^*P* ≤ 0.045).

Since AIEC abundance in human IBD patients can vary by intestinal region, we sought to determine if there are regional abundance differences of intestinal *m*AIEC in *Ptpn2*-deficient mice (Darfeuille-Michaud et al., 2004; Martinez-Medina et al., 2009; Palmela et al., 2018). KO mice show a striking increase in abundance of lumenal and mucosal-associated *m*AIEC compared with WT and Het mice (Figure 2B,C). Additionally, KO mice show increased translocation of *m*AIEC into the spleen, but not mesenteric lymph nodes, compared with WT and Het mice although this did not reach significance (Figure 2D). These data show that loss of *Ptpn2* in the mouse promotes expansion of a putative AIEC.

### Confirmation of Adherent and Invasive Properties of a Novel Mouse AIEC

Since AIEC have defined criteria to establish their adherent and invasive properties, we next determined if our novel putative *m*AIEC meets the defined criteria of an AIEC compared with the human AIEC LF82 and the non-invasive *E. coli* K12 (Martinez-Medina and Garcia-Gil, 2014). We used human Caco-2 brush border (Caco-2bbe) intestinal epithelial cells since we did not observe a difference in *m*AIEC adherence between young-adult mouse colonic intestinal epithelial cells (YAMC) and Caco-2bbe (data not shown). *m*AIEC showed greater adherence to and invasion of IECs compared with the control *E. coli* K12 (Figure 3A,B). Of note, adherence and invasion of the human LF82 AIEC to human IECs was much greater than *m*AIEC. However, *m*AIEC showed greater infection of murine Mϕ compared with human LF82 AIEC and K12 (Figure 3C). These data confirm that our novel mouse *E. coli* is phenotypically an AIEC.

**Figure 3.**
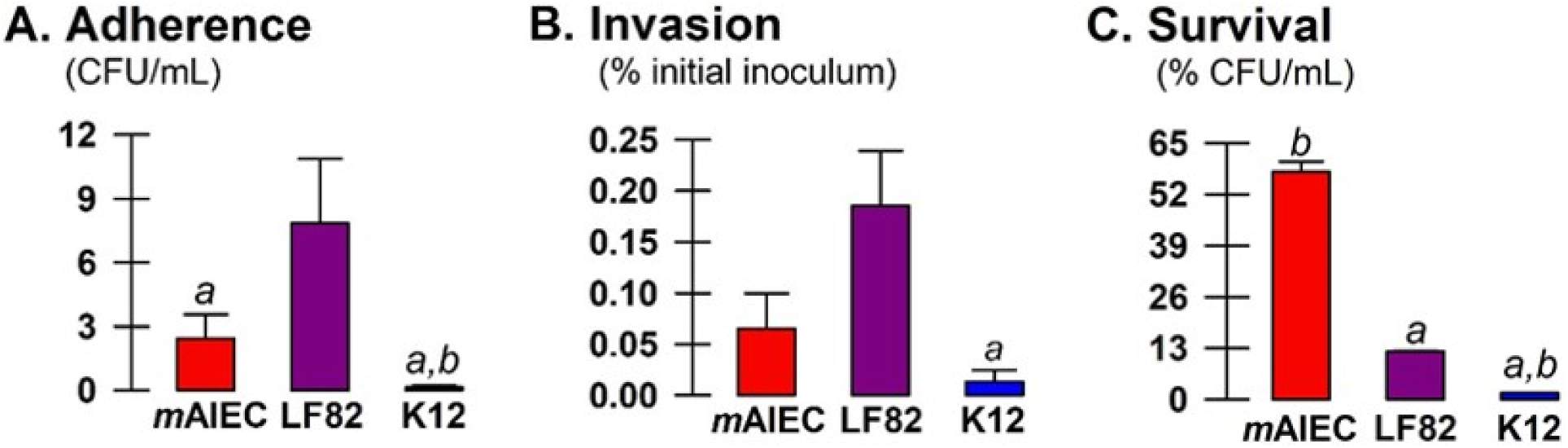
Confirmation of adherent-invasive properties of a novel mouse AIEC. Since AIEC have defined criteria to establish adherent and invasive properties, we next determined if our novel putative *m*AIEC meets the definition of an AIEC compared with the human AIEC LF82 and the non-invasive *E. coli* K12. Caco2-bbe cells (***A***,***B***) and J774A.1 murine macrophages (ATCC) (***C***) were seeded at 2 × 10^5^ cells/well in a 24-well plate until confluent and then exposed for 3 hrs to individual bacteria (MOI 10 bacteria/cell). One-way ANOVA revealed interactions for all variables (*P* ≤ 0.034; *n* = 2, 3 experiments). ***A***, Adherence, ^*a*^*P* < 0.001 *cf*. LF82, ^*b*^*P* = 0.049 *cf. m*AIEC. ***B***, Invasion, ^*a*^*P* = 0.045 *cf*. LF82. ***C***, % survival, ^*a*^*P* < 0.001 *cf. m*AIEC, ^*b*^*P* < 0.001 *cf*. LF82. These data show that *m*AIEC attach to IECs, and can replicate within macrophages, thus confirming AIEC properties.

### mAIEC Invades Germ-Free Mice

To further confirm the invasive properties of *m*AIEC, we infected germ-free (GF) mice with *m*AIEC or K12. Mono-colonization of GF mice with *m*AIEC caused sustained weight loss compared with K12 mono-colonized male GF mice, and intriguingly, *m*AIEC-dependent weight loss was restricted to male GF mice (Figure 4). Despite lack of a significant effect of *m*AIEC in female mice, mono-colonization of female GF mice with *m*AIEC caused up to 15% body weight fluctuations compared with K12 mono-colonized GF female mice, despite similar bacterial loads throughout the experiment between all the groups (data not shown). These data demonstrate that *m*AIEC is capable of causing disease in GF mice.

**Figure 4.**
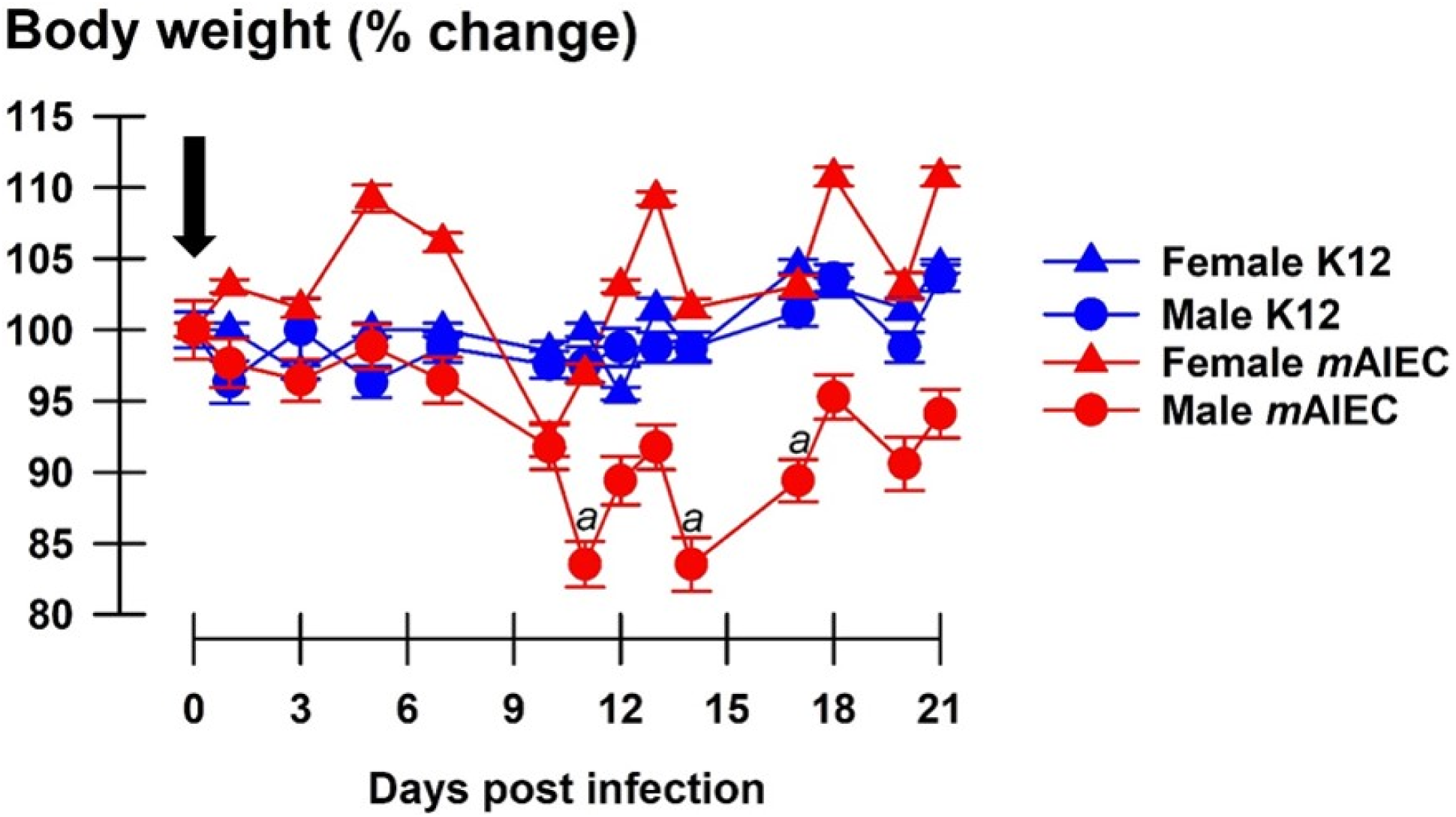
*m*AIEC colonizes and invades germ-free mice. Oral gavage of *m*AIEC (10^9^bacteria/mouse in 100µL PBS; black arrow) to germ-free mice (C57Bl/6; 12-13 weeks old) caused a selective decrease in body weight in male vs. female mice while control K12 *E. coli* was without effect after 21 days (*n* = 3). 2-way RM ANOVA revealed an interaction (*P* < 0.001). Although female *m*AIEC infected mice showed fluctuations in body weight compared with female K12-infected mice, this did not reach significance (*P* ≥ 0.41). Body weight of male *m*AIEC infected mice was significantly lower at days 11, 14, and 17 post-infection compared with male K12 infected mice (^*a*^*P* ≤ 0.02).

### mAIEC Colonizes and Increases Intestinal Permeability of SFB-free mice

We next validated that our novel *m*AIEC colonizes mouse intestine and causes disease. Whereas H_2_O-PBS and K12 mice showed stable, if not increased body weights, H_2_O-*m*AIEC mice showed significant body weight loss at 4 and 7 days post infection compared with H_2_O-PBS mice (Figure 5A). Interestingly, the drop in body weight coincided with a mild increase in disease activity index (DAI) with significantly higher DAI at day 5 and 6 post infection (Figure 5B), colon shortening (Figure 5C) and increased spleen weight (Figure 5D) 7 days post infection in H_2_O-*m*AIEC mice compared with H_2_O-PBS and K12 mice. These same mice had increased FD4 permeability (Figure 5E) on day 7 post-infection even when DAI scores had returned to normal. These data demonstrate that *m*AIEC are able to cause transient disease activity.

**Figure 5.**
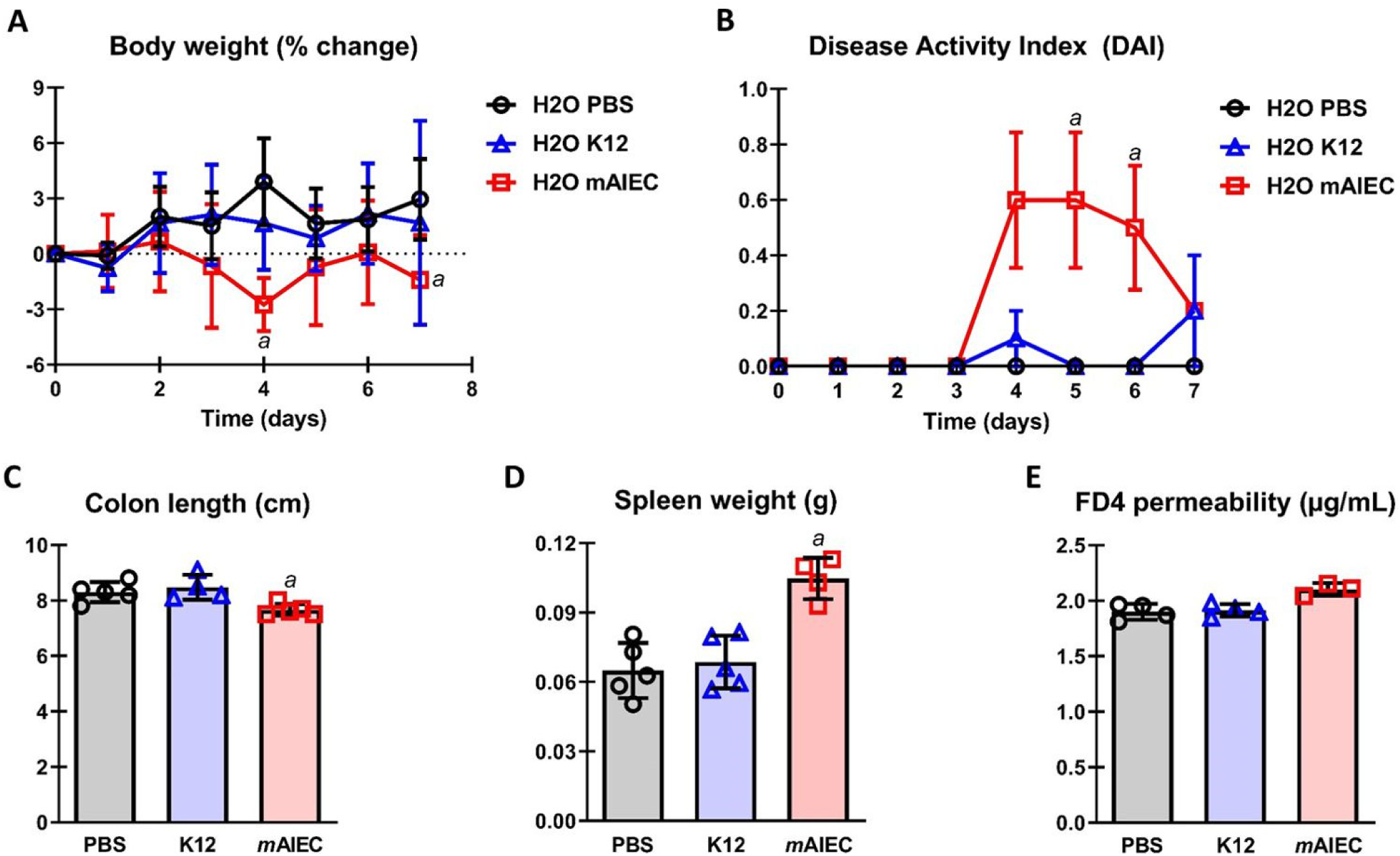
*m*AIEC colonizes and increases permeability of SFB-free mice and worsens DSS colitis. Oral gavage of *m*AIEC (10^9^ bacteria/mouse in 100µL PBS; <u>day 0-3</u>) caused **A**) weight loss (*n* = 5); **B**) mild disease (*n* = 5); **C**) colon shortening (*n* = 4-5); **D**) splenomegaly (*n* = 4-5; also see Fig 6D); and **E**) intestinal FD4 permeability (*n* = 3-4) in confirmed SFB-free C57Bl/6 mice (10 weeks old; 20-22g; JAX labs) compared with control *E. coli* (K12) or PBS after 7 days. 2-way RM ANOVA of body weight (mean ± SD) and DAI (mean ± SEM) revealed an interaction (*P* ≤ 0.002). *m*AIEC reduced body weight at day 4 and 7 compared with PBS (^*a*^*P* ≤ 0.04). One-way ANOVA of colon length, spleen weight, and FD4 permeability revealed an interaction (*P* ≤ 0.01). *m*AIEC reduced colon length, and increased spleen weight and FD4 permeability compared with PBS and K12 (^*a*^*P* ≤ 0.03).

### mAIEC worsens DSS colitis

We next wanted to determine if *m*AIEC modifies inflammation by worsening the response to an acute inflammatory episode. Mice infected with *m*AIEC and treated with DSS (DSS-*m*AIEC) mice showed a significant and sustained drop in body weight beginning at 5 days compared with DSS-PBS and K12 mice which did not show a change in body weight compared with H_2_O-PBS or K12 mice (Figure 6A). Consequently, DSS-*m*AIEC mice showed severe disease indicated by a 4 to 8-fold increase in DAI at 7 days and a further shortening in colon length compared with H_2_O-PBS, H_2_O-K12, DSS-PBS, and DSS-K12 mice (Figure 6B,C). Although DSS induced splenomegaly independent of bacterial infection (Figure 6K), DSS-*m*AIEC mice showed higher DAI as evident by increased histological score and MPO activity compared with DSS-PBS and K12 mice (Figure 6D,E). We assessed epithelial integrity and found that all mice receiving DSS showed increased permeability to FD4 compared with all H_2_O mice and this effect was independent of bacterial infection (Figure 6F). Furthermore, DSS-*m*AIEC mice showed higher levels of bacterial translocation in liver, and spleen compared with H_2_O mice and DSS-PBS and K12 mice (Figure 6G,H); however, there was no significant translocation of bacteria found in distal colon or MLN in all groups (Figure 6I,J). Overall, these data confirm that *m*AIEC: (i) invades SFB-free mouse intestine, (ii) causes disease and weight loss; (iii) increases intestinal permeability; and (iv) exacerbates disease in the setting of inflammation.

**Figure 6.**
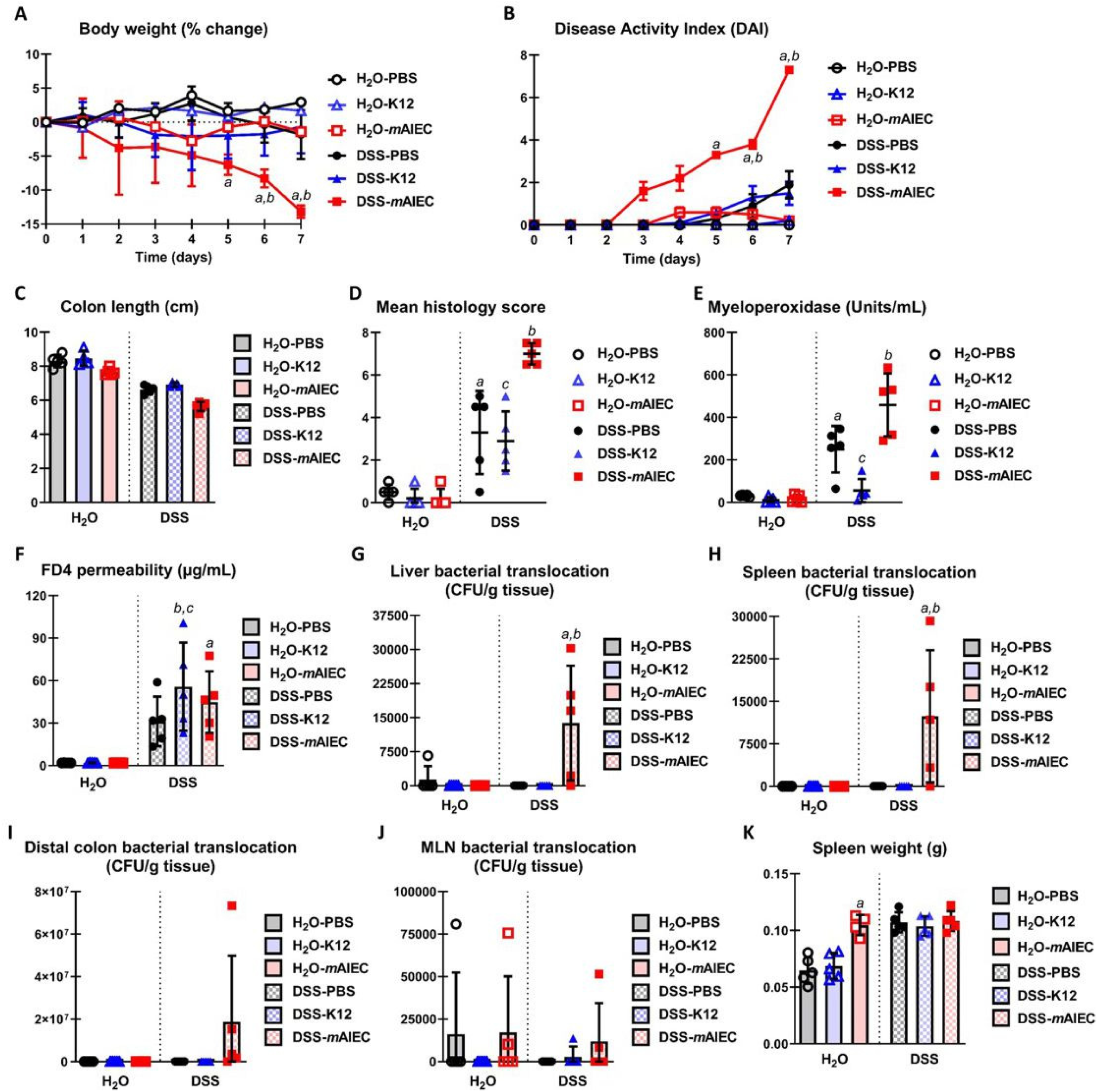
*m*AIEC worsens DSS colitis. Co-administration of *m*AIEC and DSS (2.5%; 7 days) in drinking water exacerbated **A**) weight loss; **B**) severe disease; **C**) colon shortening; **D**) mean histology score; **E**) MPO activity; **F**) FD4 permeability; and translocation of *m*AIEC in the **G**) liver; and **H**) spleen; but not **I**) distal colon or **J**) MLN and no additional increase in **K**) spleen weight in confirmed SFB-free C57Bl/6 mice (10 weeks old; 20-22g; JAX labs) compared with control *E. coli* (K12) or PBS. 2-way RM ANOVA of body weight and 1-way ANOVA of DAI revealed an interaction (*P* < 0.001). *m*AIEC potentiated the loss in body weight (5-7 days ^*a*^*P* ≤ 0.01 *cf*. DSS-PBS and 6-7 days ^*b*^*P* ≤ 0.05 *cf*. K12), and increase in DAI (mean ± SEM, 7 days ^*a*^*P* ≤ 0.005 *cf*. all groups). Two-way ANOVA of all other parameters revealed an interaction (*P* ≤ 0.002) except with distal colon length, and distal colon and MLN translocation (*P* ≥ 0.15). DSS-PBS mice showed an increase in mean histology score and MPO activity (^*a*^*P* ≤ 0.005 *cf*. H_2_O mice) and this effect was worsened when DSS mice were infected with *m*AIEC (^*b*^*P* ≤ 0.005 *cf*. DSS-PBS). Mean histology score was not increased in DSS-K12 mice (^*c*^*P* = 0.99 *cf*. DSS-PBS). MPO activity was lower in DSS-K12 mice (^*c*^*P* = 0.01 *cf*. DSS-PBS). DSS-*m*AIEC mice showed increased FD4 permeability (^*a*^*P* = 0.01 *cf*. H_2_O mice); however, DSS-K12 mice also showed increased permeability to FD4 (^*b*^*P* = 0.01 *cf*. H_2_O mice) with no difference with DSS-mAIEC mice (^*c*^*P* = 0.92 *cf*. DSS-K12). DSS-*m*AIEC mice had greater translocation of bacteria in their liver and spleen (^*a,b*^*P* ≤ 0.06 *cf*. H_2_O mice and DSS-PBS and K12 mice).

### mAIEC prevents recovery from DSS colitis

Since *m*AIEC worsened acute colitis, we wanted to determine if it also impaired recovery from colitis. DSS induced colitis in all of the mice in the recovery group regardless of bacterial infection as evident by reduced body weight; however, rec-*m*AIEC mice showed a more severe body weight loss and slower recovery compared with rec-PBS and rec-K12 mice (Figure 7A). Consequently, rec-*m*AIEC mice showed a higher and more sustained DAI from day 8 through 14 compared with rec-PBS and K12 mice (Figure 7B). This coincided with splenomegaly and a decrease in colon length in rec-*m*AIEC mice compared with rec-PBS and K12 mice (Figure 7C), however no significant increase in translocation of bacteria in distal colon was observed (Figure 7D). Overall, these data show that *m*AIEC prevents recovery of acute colitis.

**Figure 7.**
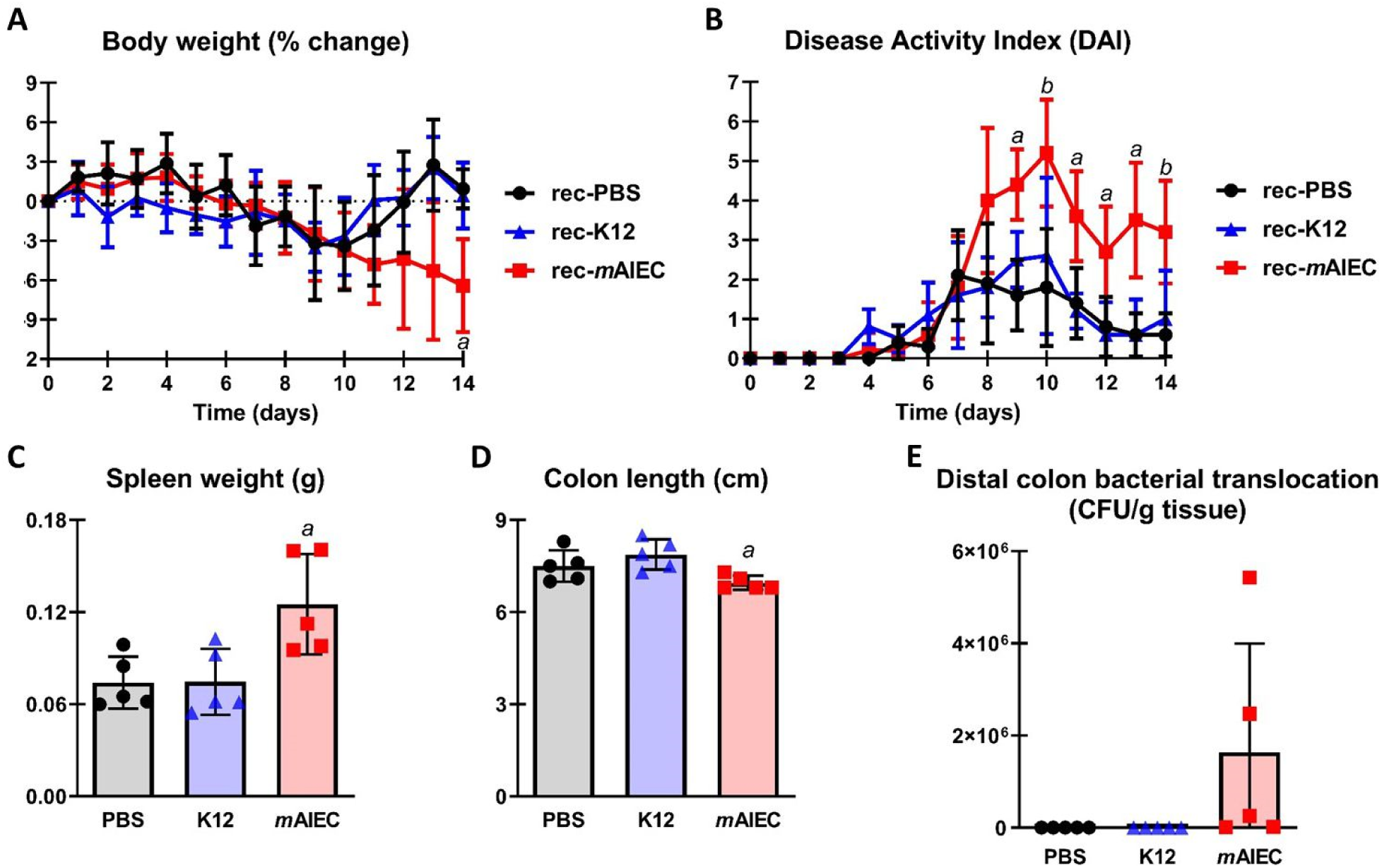
*m*AIEC impairs recovery from colitis. Administration of *m*AIEC post DSS (2.5%; 7 days) in drinking water impaired recovery of **A**) body weight; **B**) disease; **C**) splenomegaly; and **D**) colon length with no effect on **E**) translocation of *m*AIEC to the distal colon in confirmed SFB-free C57Bl/6 mice (10 weeks old; 20-22g; JAX labs) compared with control *E. coli* (K12) or PBS. Two-way RM ANOVA of body weight and DAI revealed an interaction (*P* < 0.001). *m*AIEC prevented against recovery of body weight loss (rec-*m*AIEC 14 days ^*a*^*P* ≤ 0.02 *cf*. rec-PBS and K12), and decrease in DAI (rec-*m*AIEC 9, 11, 12, and 13 days ^*a*^*P* ≤ 0.04 *cf*. rec-PBS and K12 and 10 and 14 days ^*b*^*P* ≤ 0.02 *cf*. rec-PBS only). One-way ANOVA of spleen weight revealed an interaction (*P* = 0.009). *m*AIEC increased spleen weight during the recovery phase (rec-*m*AIEC ^*a*^*P* ≤ 0.02 *cf*. rec-PBS and K12). No effect of translocation of bacteria in the recovery groups (*P* = 0.13).

### Genome sequence analysis of mAIEC

Since the *m*AIEC isolate showed 100% sequence identity and match of the ITS region to the human AIEC LF82 (see Fig 1A), to identify how similar these bacteria are across their complete genomes, we sequenced the genome of our novel mouse AIEC and compared it with the published genome sequence of LF82 (GenBank CU651637.1). *m*AIEC showed approximately 90.3% similarity to the human AIEC LF82. We further probed for the presence of putative attachment genes found in *E. coli’s* and other AIECs including LF82 in our *m*AIEC. Attachment genes common between *m*AIEC, another non-pathogenic mouse AIEC NC101, and the human AIEC LF82, or unique to each isolate, were compared by BLAST analysis (Tables 1). There were 14 total attachment genes that were used in this analysis. In particular, *m*AIEC showed 3 distinct differences in attachment genes compared with LF82 and NC101: 1) *yadC* gene had the lowest % similarity, 2) *papA* gene was strictly unique to *m*AIEC, and 3) *pic* gene was unique to mouse AIEC (Table 1). There were no attachment gene overlap between *m*AIEC and LF82 or between LF82 and NC101 (Figure 8).

**Table 1.** Summary of Adherence Genes Alignment Between *m*AIEC and the LF82 and NC101 AIECs.

| Virulent Gene | Reference | Function | % Identity with NC101 | % Identity with LF82 (Accession CU651637.1) |
| --- | --- | --- | --- | --- |
| *fimH* | (Barnich et al., 2007) | Terminal subunit of Type 1 pilus | 99.61 | 99.20 |
| *papC* | (Yang et al., 2017) | Outer membrane usher protein | 99.61 | 99.26 |
| *yadC* | (Verma et al., 2016) | Fimbrial-like protein | 85.1 | 85.95 |
| *ygi* | (Bachmann et al., 2015) | Putative acid-amine ligase | 97.80 | 99.18 |
| *csgE* | (Nash et al., 2010) | Curli assembly protein; bacterial aggregation | 99.29 | 98.91 |
| *yehA* | (Ravan and Amandadi, 2015) | Putative fimbrial-like protein | 99.35 | 99.43 |
| *hcpA* | (Nash et al., 2010) | Major exported protein | 98.36 | 100 |
| *fimA* | (Nash et al., 2010) | Type-1 fimbrial protein A chain | 98.82 | 98.56 |
| *papA* | (Feria et al., 2001) | Pap fimbrial major pilin protein | Not expressed | Not expressed |
| *ftsh* | (Allen et al., 2008) | ATP-dependent zinc metalloprotease | 99.29 | 97.90 |
| *pic* | (Camprubi-Font et al., 2018) | Serine protease pic autotransporter | 95.21 | Not expressed |
| *malX* | (Nash et al., 2010) | PTS system maltose-specific EIICB component | 97.91 | 97.93 |
| *dsbA* | (Nash et al., 2010) | Thiol:disulfide interchange protein | 100 | 100 |
| *fliA* | (Claret et al., 2007) | Motility sigma factor | 99.62 | 100 |

**Figure 8.**
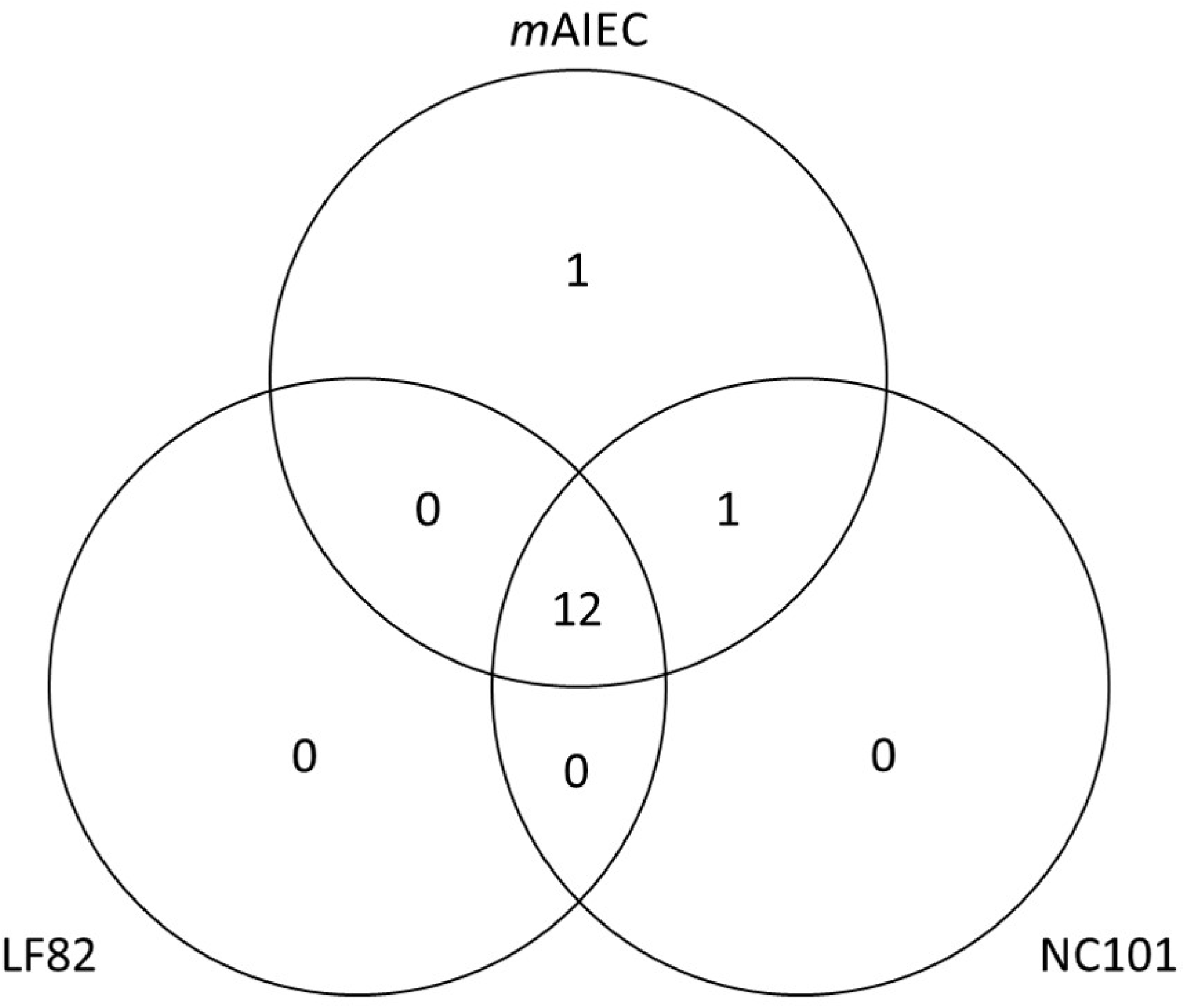
Genome of *m*AIEC displays differential adherence gene properties. Sequence alignment of 14 putative virulence genes identified common or unique attachment genes in *m*AIEC, LF82, and NC101. Identified attachment genes from each bacterial genome are represented in a Venn diagram as number of genes common or unique to each bacteria.

## Discussion

One of the factors considered a contributing factor to IBD development and progression is alteration of composition of the intestinal microbiome that favors expansion of commensal bacteria with pathogenic potential. In mice with reduced or lack of expression of the IBD risk gene, PTPN2, we found significant alterations in intestinal microbiome variance compared with wild-type littermates. Moreover, *Ptpn2*-deficient mice showed a selective increase in abundance of *Proteobacteria*, specifically *E. coli*. Intriguingly, the highest increase in abundance of lumenal and mucosal associated bacteria in *Ptpn2*-KO mice was of an *E. coli* that shared >90% genome sequence similarity to the human IBD-associated AIEC LF82. Furthermore, IBD patients that have been genotyped for the *PTPN2* IBD risk allele rs1893217 show altered microbiomes and increased presence of *Proteobacteria* thus further validating our in vivo system as a model of IBD (Yilmaz et al., 2018; Martinez-Medina and Garcia-Gil, 2014; Knights et al., 2014). Although LF82 was originally isolated from the ileum of a Crohn’s disease patient, additional AIEC have since been identified in ulcerative colitis patients and isolated from other intestinal regions (Darfeuille-Michaud et al., 1998; O’Brien et al., 2016; Lee et al., 2019; Kotlowski et al., 2007). Whereas *m*AIEC was detected albeit at low levels in lumenal and mucosal-associated samples from *Ptpn2* wild-type mice indicating that this bacterium is part of the resident microbiota, *Ptpn2*-KO mice showed higher abundance of *m*AIEC both in the lumen and associated with epithelia in small and large intestinal regions of the gut. Overall, reduced expression of the IBD candidate gene, *PTPN2*, alters intestinal microbial communities in mice to favor expansion of a novel mouse AIEC.

We confirmed the adherence, invasive, and survival properties of *m*AIEC to the well-studied human AIEC LF82 (Martinez-Medina and Garcia-Gil, 2014). We found that *m*AIEC adhere to and invade intestinal epithelial cells, although to a lesser extent than LF82. Although adherence and invasion of *m*AIEC was lower than LF82, this was expected as we were using human IECs with a mouse bacterium. However, the absolute values of *m*AIEC adherence and invasion are not significantly different from the published adherence and invasive properties for an AIEC, and there was no difference in adherence of *m*AIEC to young-adult mouse colonic cells compared with human IECs (Martinez-Medina and Garcia-Gil, 2014; Darfeuille-Michaud et al., 2004). Intriguingly, survival of *m*AIEC in murine macrophages was significantly higher than LF82, and both *m*AIEC and LF82 were significantly higher than K12. Given that these studies were performed with mouse macrophages (J774.1) this may suggest that our novel *m*AIEC represents a better tool to study AIEC induced disease in a mouse host than the human isolate, LF82. Overall, these data confirm the epithelial adherent and invasive properties and survival in macrophages of our novel *m*AIEC compared with established parameters.

We first validated whether our novel mouse AIEC can cause disease in mice. Indeed, we found that mono-colonization with *m*AIEC reduced body weights of germ-free mice. Interestingly, this affect seemed to be specific to male mice; however, female mice also showed large fluctuations in body weight indicating that *m*AIEC resulted in a response in both male and female germ-free mice. Although we found that *m*AIEC can invade germ-free mice, this setting does not reflect the conditions of which we found high abundance of *m*AIEC (SPF mice with low abundance of SFB, data not shown). Therefore, we infected mice harboring a microbiome (SPF mice). We confirmed that mono-infection of *m*AIEC but not K12 results in mild disease evident by reduced body weight and slightly increased disease activity. The lack of an overt response to infection may be explained by the fact that these mice already have a microbiome that may be partially outcompeting or countering *m*AIEC pathogenicity. Never-the-less, we showed for the first time that the mouse resident bacterium (pathobiont) *m*AIEC can cause transient disease under normal conditions without additional stressors.

Since we isolated *m*AIEC from *Ptpn2*-KO mice that develop systemic inflammation and since AIEC have been shown to be increased in CD and UC patients, we investigated the effects of *m*AIEC in an inflammatory setting. Mice exposed to mild DSS to induce colitis and simultaneously infected with *m*AIEC showed a more severe disease (greater body weight loss, increased disease scores, bacterial translocation) and increased macromolecular permeability. In fact, AIEC have been shown to facilitate a response to a flare in IBD patients and ability of an IBD-susceptible host to recover from an ‘event’ such as acute colitis caused by ingestion of a foodborne pathogen thus potentiating IBD progression (Mirsepasi-Lauridsen et al., 2019). Of note, mice infected with *m*AIEC fail to recover (body weight loss and increased disease activity) from DSS-induced colitis (Small et al., 2016).

We compared the genome sequence of our mouse AIEC to the human AIEC LF82 and a non-pathogenic mouse AIEC, NC101. *m*AIEC showed >90% similarity to LF82, indicating its similarity to LF82 yet that it’s distinct from LF82. Interestingly, *m*AIEC contained a distinct putative attachment gene, *papA*, that was not found in LF82 or NC101. Additionally, we identified a putative attachment gene, *pic*, that was only expressed in mouse AIEC that may be used to distinguish between mouse and human AIEC. Overall, we identified distinct putative attachment genes in *m*AIEC thereby also demonstrating the novelty of this AIEC.

While a number of studies have investigated potential contributions of commensal bacteria that possess or acquire pathogenic potential, such as the LF82 AIEC in the pathogenesis of IBD, it still remains elusive as to how IBD susceptibility genes modulate the intestinal microbiome to maintain intestinal homeostasis and how disruption of this interaction precipitates changes in the microbiome that promote disease. We show here for the first time in an untreated mouse model of IBD (*Ptpn2*-deficient), how loss of an IBD associated gene, *PTPN2*, alter the intestinal microbiome and favors expansion of a novel mouse AIEC that is capable of both initiating and exacerbating disease. Thus, PTPN2 plays a key role as a “microbial modulator” of the microbiome to protect against pathobiont expansion and colonization. We propose that *Ptpn2*-deficient mice may serve as a useful model to investigate how host genetics modulate the balance of intestinal microbes, and that this novel mouse AIEC can be utilized to interrogate mechanisms of pathobiont-induced intestinal inflammation.

